# Mouse pups lacking a cerebral cortex develop abnormal vocal behavior

**DOI:** 10.1101/2025.07.17.665371

**Authors:** J. Lomax Boyd, César D. M. Vargas, Erich D. Jarvis

## Abstract

Vocal learning is an essential component of spoken language and critically depends on the cerebral cortex. The evolutionary origins of cortical/pallial control over vocal learning abilities in mammals and songbirds remains largely unknown. For instance, reports conflict on whether the cerebral cortex contributes, in any way, to vocal communication in vocal non-learning mice. Physiological studies in adult mice have shown that regions of the motor cortex have roles in modulation of vocalizations in mice, yet, genetic ablation of the cerebral cortex reportedly has minimal, if any, impact on mouse vocal behavior. Re-analysis of adult acortical mice revealed that deep learning machine classifiers could distinguish mutant ultrasonic vocalizations from wildtypes. However, the specific acoustic features underlying these differences were not identified. Here, we investigated isolation calls of acortical mouse pups using statistical analysis of acoustic features and playback experiments to determine whether mutants lacking a cerebral cortex have altered vocal development. We find that a subset of acoustic features differ between acortical and wildtype pup vocalizations and that these differences are indicative of distress. Moreover, call bouts of acortical pups have lower informational complexity that are more comparable to random probability sampling. Playbacks indicate that dams preferentially approach vocalizations of acortical pups. Our analyses provide evidence that the murine cerebral cortex influences development of complex vocal behaviors, suggesting mice can be used to gain useful insights into the foundations of vocal learning.

## Introduction

The cerebral cortex plays an essential role in the evolution, development, and dysregulation of human speech and language. Among the components that contribute to the complexity of spoken language–which include, auditory learning, semantics, syntax, and speech perception– vocal learning represents a critical yet rare ability to voluntarily modulate vocal production based on social experience and auditory feedback. This component of human speech evolved independently in five mammalian and three avian lineages^1^. Changes in cortical/pallial size, organization, and connectivity have been hypothesized as the neuroanatomical basis of vocal learning evolution, yet it remains challenging to experimentally investigate these hypotheses in mammalian vocal learners. Jarvis and colleagues (2012) proposed that variation in vocal learning abilities can be conceptualized along a continuum, with many species possessing the ability to learn sound associations (auditory learning) or produce innate vocalizations in new contexts (vocal usage learning), while significantly fewer species being capable of producing novel vocalizations based on experience (vocal production learning)^2^. Moreover, others have extended the continuum hypothesis of vocal learning into multiple dimensions, with each axis representing modules of temporal and acoustic complexity whose neurobiological and evolutionary inter-dependence remains an open question^3^. For instance, all vocal learning species examined to date possess a specialized, robust motor cortex-to-brainstem motor neuron direct connection that helps to control phonatory muscles (mammal larynx; avian syrinx) and vocalization^1,4^. The discovery that mice, a vocal non-learning species, have a similar, yet sparse connection from the primary motor cortex (M1) to laryngeal motor neurons demonstrated that the neural basis of vocal learning has deep evolutionary and anatomical origins in the mammalian lineage, which can vary in magnitude across species with differing vocal learning abilities. Furthermore, this discovery suggested that cortical vocal-motor circuits could be investigated in an experimentally tractable model system^5^.

Various experiments in adult mice have shown that the cortex has some role in controlling vocal behavior and vocal musculature. For example, lesions to a laryngeal area of M1 were found to impact the ability of mice to control the frequency distribution of vocalizations emitted during courtship interactions^5^. Recent electrophysiological work showed that stimulation of mouse M1 can produce laryngeal muscle contractions with latencies of stimulation-induced contractions consistent with a sparse indirect projection reported earlier from the motor cortex to phonatory motor neurons^6^. Comparative work in another rodent species, Alston’s singing mice (*Scotinomys teguina*), also showed that the orofacial motor cortex in M1 is involved in the temporal control of vocal flexibility used during social turn-taking^8,9^. Despite these physiological results suggesting mouse M1 may have a role in controlling mouse vocal musculature and behavior, a study investigating the vocal behavior of *Esco2^fl/fl^;Emx1-Cre* mice, which are missing a majority of cortical areas (acortical), reported few, if any, changes in vocal behavior compared to wildtype control mice^10^. The Hammerschmidt et al. (2015) study suggested the cortex in mice has little control over vocal behavior, courtship behaviors, or reproduction^11,12^. Using deep neural networks, Ivanenko et al. (2020) re-analyzed spectrograms of the acortical adult vocalizations and found that machine classifiers could distinguish acortical from wildtype vocalizations with accuracies (~63.4%) above those expected by chance. The study, however, did not re-analyze vocal behavior of post-natal day 9 mouse pups (P9) originally reported in Hammerschmidt et al. (2015).

To clarify the role of the cortex in development of vocal behaviors, we re-analyzed vocalizations from acortical mouse pups, which were generously provided by Kurt Hammerschmidt and originally reported in Hammerschmidt et al (2015). Using a K-means classifier, analysis of acoustic/spectral/temporal features, and ultrasonic playback experiments with maternal dams caring for age-match litters, we discovered that the vocal behavior of acortical mouse pups are distinguishable from those of wildtype animals, and that these changes influence maternal dam behavior. We report evidence that the vocal repertoire of acortical pups is acoustically distinct from wild types with significant reductions in informational content that implicate the cortex in regulation of acoustic-temporal complexity.

## Results

### Production of maternal isolation calls

Maternal-isolation induced calls of *Esco2^+/+^;Emx1-Cre* (n=8), wildtype (WT), and acortical *Esco2^fl/fl^;Emx1-Cre* (n=10), knockout (KO), mouse pups at post-natal day 9 (P9) were re-analyzed^10^. Ultrasonic vocalizations (USVs) were detected and vocal features (**Table 1**) extracted using MSA2, a Python implementation of Mouse Song Analyzer (MSA)^5,13–15^. Overall, vocal production rates were similar between WT and acortical KO mice (Shapiro-Wilk normality test: p=0.0023, W=0.813; Wilcoxon rank sum test: p=0.897, W=42; **Figure 1A**). However, MSA2 detected more individual USVs compared to previous reports^10^. We classified USVs into five syllable-type groups based on the presence of pitch jumps: simple (s) syllables with no pitch jump, down-jump (d), up-jump (u), multi-jump (M), and unclassified (UC) USVs^10^. Manual inspection of spectrograms revealed unclassified syllables were primarily composed of USVs with pitch jumps and durations that met the minimum threshold of detection (6 ms). On average, more than half (51.3%) of USVs emitted by pups were simple ‘s’ syllables with no overall difference between WT and acortical pups (ANOVA: p=1, df=80, F=0). However, significant genotype-by-syllable type interactions were detected (ANOVA: p=0.00772, df=80, F=3.74) with KOs producing 63.4% more up-jump syllables (Tukey post hoc: p=0.0026, df=80, t=-3.11) and 37.3% fewer unclassified syllables, compared to controls (Tukey post hoc: p=0.0318, df=80, t=2.19; **Figure 1B**).

**FIGURE 1.**
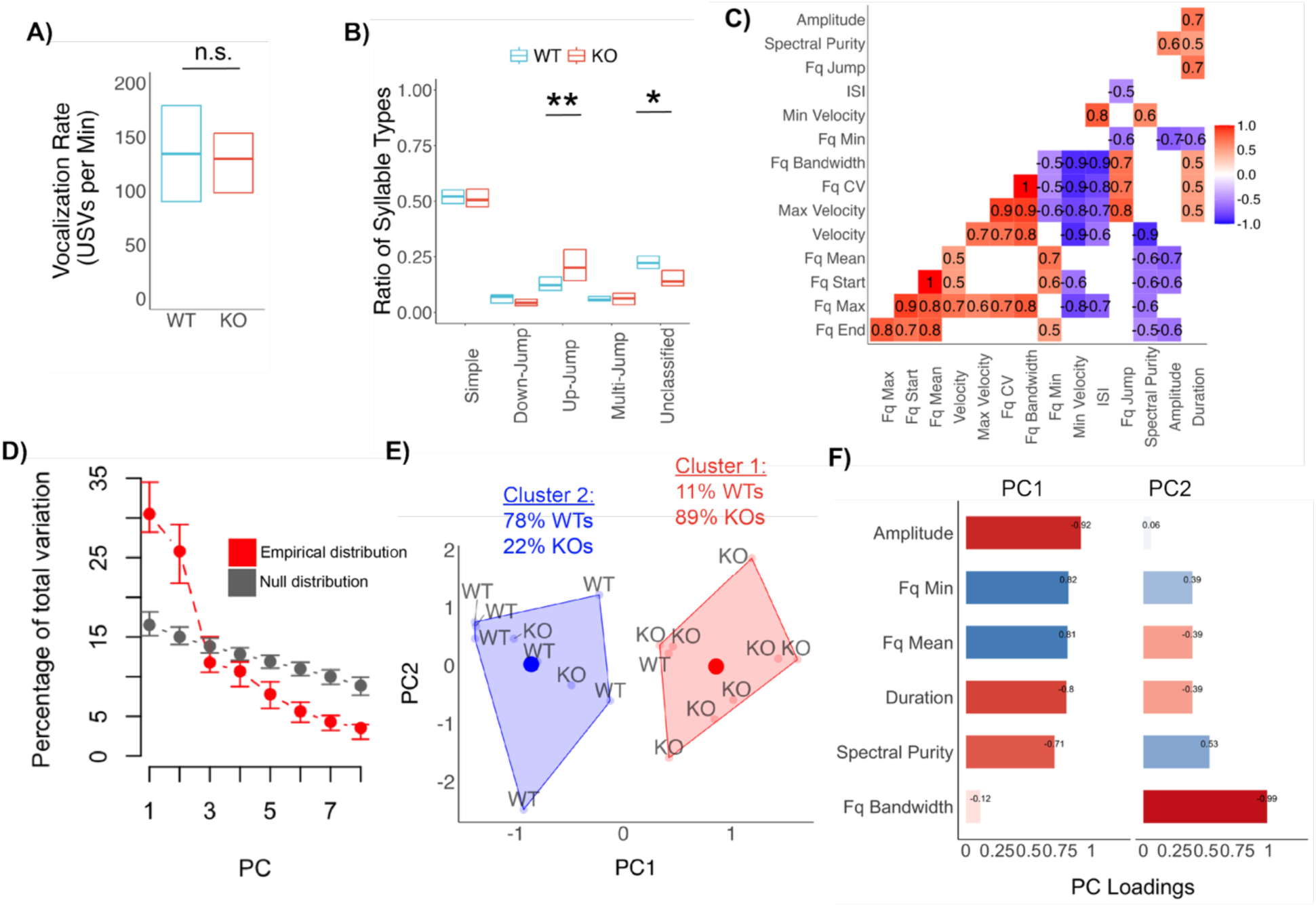
Acortical mouse pups develop distinct vocal repertoires. Ultrasonic vocalizations (USV) of isolation-induced calls from *Esco2^+/+^;Emx1-Cre* (wildtype, WT) and *Esco2^fl/fl^;Emx1-Cre* (knockout, KO) pups were re-analyzed from Hammerschmidt et al. 2015^10^. **A**) Vocalization rates were similar between acortical mutants (red; median, 129.6.7±85 USVs per minute) and WTs (blue; median, 134.3±57 per minute) mouse pups (p=0.897, W=42). **B**) The proportion of USVs classified into five syllable types based on the presence of pitch jumps. Acortical KOs mice produced significantly more up-jump syllables (p=0.0026, t=-3.11, df=80) and fewer unclassified syllables (p=0.0318, t=2.19, df=80), compared to controls. Boxplots represent the upper 75^th^ and lower 25^th^ quartiles of data with median values visualized by thicker horizontal line. **C**) Pearson correlation matrix of vocal features with significant positive (red) and negative (blue) correlations. Non-significant correlations (p>0.05) are blank. **D**) Comparison of variation accounted for by each principal component (PC) in the empirical data and a null distribution of randomly re-sample data with confidence intervals generated using 1,000 bootstrap replicates. **E**) K-means clustering performed on PCA scores from the first two PCs for vocal features. **F**) Vocal feature loadings for PC1 and PC2. Production of syllable types analyzed using linear mixed models with repeated measures for inter-individual correlations across pups with Tukey adjustment for multiple test correction. Abbreviations: ‘Fq’, frequency; ‘CV’, coefficient of variance, ‘ISI’, inter-syllable interval, ‘n.s., not significant; ‘*’, p≤0.05; ‘**’, p≤0.01.

**Table 1.**
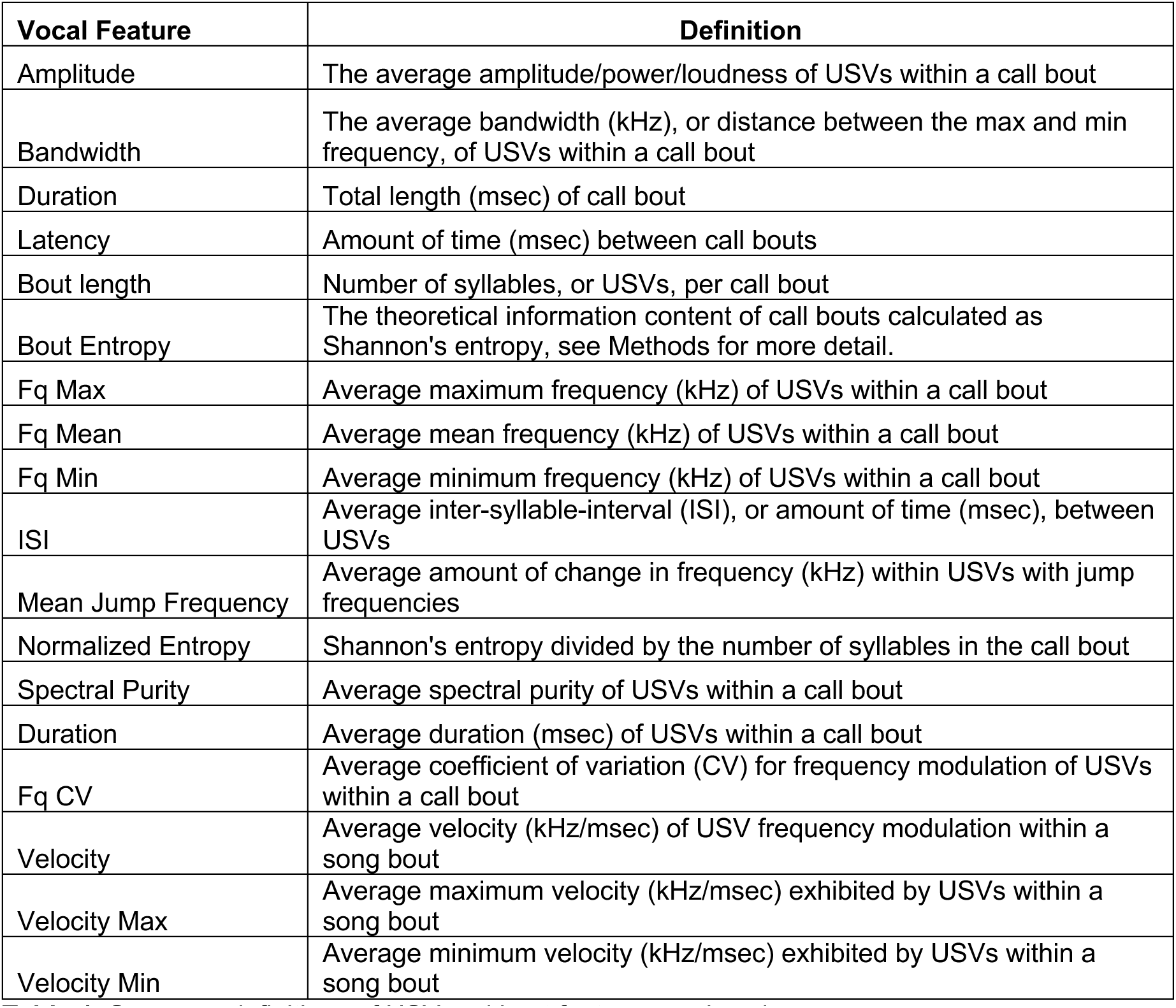
Summary definitions of USV and bout features analyzed.

| Vocal Feature | Definition |
| --- | --- |
| Amplitude | The average amplitude/power/loudness of USVs within a call bout |
| Bandwidth | The average bandwidth (kHz), or distance between the max and min frequency, of USVs within a call bout |
| Duration | Total length (msec) of call bout |
| Latency | Amount of time (msec) between call bouts |
| Bout length | Number of syllables, or USVs, per call bout |
| Bout Entropy | The theoretical information content of call bouts calculated as Shannon's entropy, see Methods for more detail. |
| Fq Max | Average maximum frequency (kHz) of USVs within a call bout |
| Fq Mean | Average mean frequency (kHz) of USVs within a call bout |
| Fq Min | Average minimum frequency (kHz) of USVs within a call bout |
| ISI | Average inter-syllable-interval (ISI), or amount of time (msec), between USVs |
| Mean Jump Frequency | Average amount of change in frequency (kHz) within USVs with jump frequencies |
| Normalized Entropy | Shannon's entropy divided by the number of syllables in the call bout |
| Spectral Purity | Average spectral purity of USVs within a call bout |
| Duration | Average duration (msec) of USVs within a call bout |
| Fq CV | Average coefficient of variation (CV) for frequency modulation of USVs within a call bout |
| Velocity | Average velocity (kHz/msec) of USV frequency modulation within a song bout |
| Velocity Max | Average maximum velocity (kHz/msec) exhibited by USVs within a song bout |
| Velocity Min | Average minimum velocity (kHz/msec) exhibited by USVs within a song bout |

### Characterizing the vocal repertoire of pup calls

We next sought to investigate whether acortical mouse pups exhibited global differences in the acoustic/spectral/temporal characteristics of USVs, compared to wildtype animals. First, we examined the correlational structure of vocal features to identify highly correlated factors that measure similar vocal features of USVs (**Figure 1C**). For instance, frequency bandwidth and coefficient of variance (CV) are nearly perfectly correlated (corr=1), and thus, represent redundant measures of frequency modulation. Using a subset of vocal features with correlations less than 0.80 (mean frequency, minimum frequency, amplitude, spectral purity duration, and bandwidth), we performed principal component analysis (PCA) and found PC1 and PC2 account for 83.8% (PC1: 55.3%; PC2: 28.5%, respectively) of the total variance in the data (**Supplementary Figure 1A**). We then compared the percent of variance explained by each PC for the empirical data to a null distribution of randomly re-sampled data (n=1,000 bootstrapped replicates). The percent of variance in empirical data explained by PC1 and PC2 was notably higher compared to the null distribution, which later converged with additional PCs (**Figure 1D**). Thus, PC1 and PC2 represent meaningful features of the empirical data above the null expectation. Moreover, PC1/PC2 scores were found suitable for clustering (Hopkin’s statistic, H=0.30). Next, we performed a cluster optimization procedure on PC1/PC2 scores and identified two as the optimal number of clusters based on the results of 30 cluster optimization algorithms (**Supplementary Figure 1B**). We then performed k-means clustering on PC1/PC2 scores. The goodness of classification was computed using linear discriminant analysis and found to be 88.9% for each cluster group. Acortical pups (8 of 10) were primarily assigned to Cluster 1 (89% of all cluster 1 assignments), whereas wildtype pups (7 of 8) were primarily assigned to Cluster 2 (77.8% of all cluster 2 assignments) (**Figure 1E**). PC loadings revealed that USV amplitude, minimum frequency, mean frequency, USV duration, and spectral purity were highly correlated with PC1, whereas frequency bandwidth primarily loaded unto PC2 (**Figure 1F**). These findings indicate that the vocal repertoire of acortical mouse pups are highly distinguishable from wildtype mice based on a combination of acoustic features represented along the first principal component.

### Vocal features that distinguish acortical pup vocalizations

Individual vocal features contributing to PC1 and PC2 were individually analyzed for genotypic differences between WT and acortical KO pups. Compared to controls, the vocalizations of acortical mouse pups showed significant decreases in mean frequency (p=0.0150, df=15.9, t=2.72) and minimum frequency (p=0.0189, df=15.8, t=2.62 (**Figure 2A,B**), and significant increases in amplitude (p=0.00527, df=15.8, t=-3.23), spectral purity (p=0.0462, df=15.9, t=-2.16), and duration (p=0.0234, df=15.5, t=-2.51) (**Figure 2C-E**). These vocal features loaded heavily onto PC1. The average bandwidth of USVs, which loaded heavily onto PC2, were similar (p=0.895, df=13, t=-0.132) between genotypes (**Figure 2F**).

**FIGURE 2.**
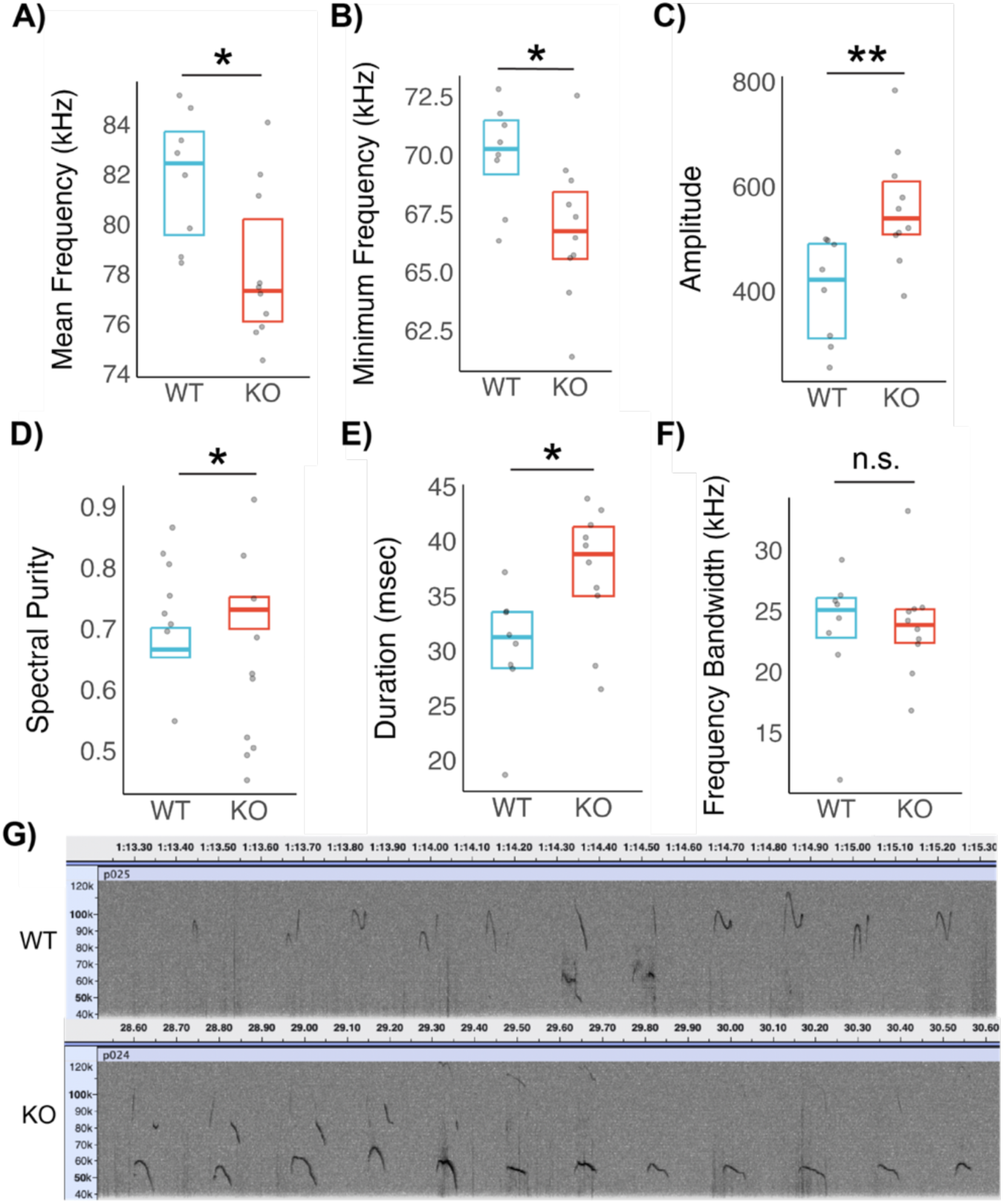
Acoustic features of acortical mouse pups. Mutant USVs displayed significant decreases in (**A**) mean frequency (p=0.015, df=15.9, t=2.72), (**B**) minimum frequency (p=0.0189, df=15.8, t=2.62), and increases in (**C**) amplitude/loudness/power (p=0.00527, df=25.8, t=-3.23), (**D**) spectral purity (p=0.0463, df=15.8, t=-2.16), and (**E**) duration (p=0.0234, df=15.5, t=-2.51), compared to control pups. **F**) Bandwidth of USVs were similar between genotypes (p=0.896, df=13, t=-0.133). Boxplots visualize the upper 75^th^ and lower 25^th^ quartiles of data with median values represented by horizonal bars. Statistical analysis of vocal features computed using Welch two-sample t-test. **G**) Representative spectrogram of WT and acortical KO USVs on a 40-120 kHz scale emitted within a 2 second time-interval. ‘n.s.’, not significant; ‘*’, p≤0.05; ‘**’, p≤0.01.

### Temporal and informational features of isolation-induced call bouts

We next investigated the temporal characteristics of acortical and wildtype pup call bouts by organizing sequences of 5 or more USVs, separated by less than 250 ms of silence (inter-syllable intervals, ISIs<250 ms), into isolation-induced call bouts. The average number of call bouts emitted by WT (28±14.4 bouts) and KO (30.1±12.6 bouts) pups were similar (Wilcoxon rank test, p=0.894, W=38) (**Figure 3A**). The average time between USVs within call bouts were also similar (p=0.695, W=34,386). However, acortical KOs exhibited significantly more variation in intra-bout ISIs (p=0.0395, W=37,252) (**Figure 3B,C**). Isolation call bouts of acortical pups were also significantly longer compared to controls (p=6.53×10^−5^, W=25,904) (**Figure 3D**).

**FIGURE 3.**
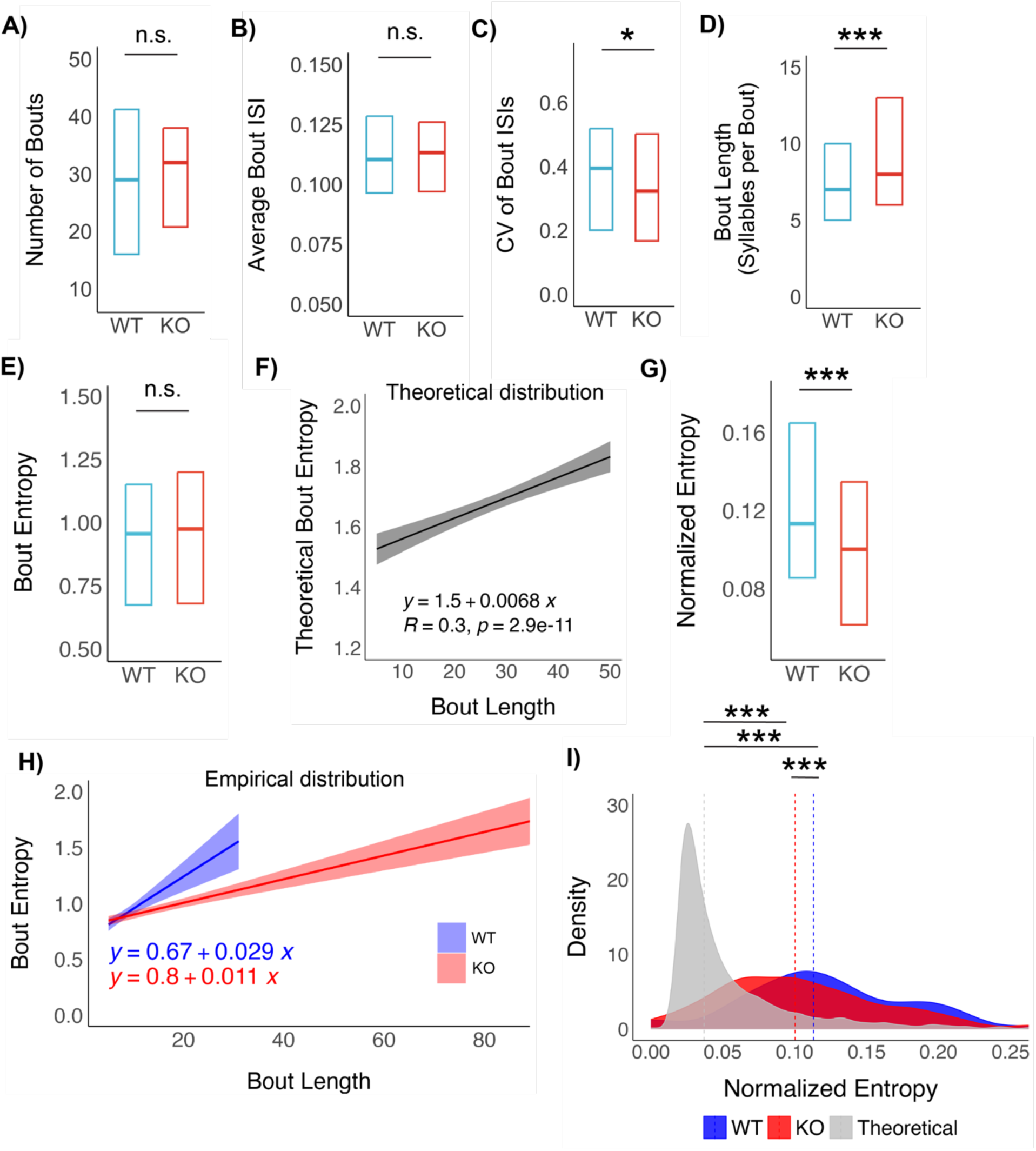
Isolation calls of acortical mouse pups have reduced temporal variation and informational complexity. **A**) Total number of call bouts were similar across genotypes (p=0.894, W=38). **B**) The average time interval between USVs in a call bout, or inter-syllable interval (ISI), were also similar (p=0.695, W=34,386). **C**) Coefficient of variation (CV) of ISIs within a bout, a standardized measure of variability, were significantly lower in KO pups, compared to WT controls (p=0.0395, W=37,252). **D**) The number of syllables per bout, or bout length, were significantly higher in acortical KOs compared to controls (p=6.53×10^−5^, W=26,904). **E**) The informational complexity of song bouts, computed as Shannon’s entropy, were similar across genotypes (p=0.530, W=32,633). **F**) The theoretical distribution of bout entropy obtained from weighted random re-sampling of syllable types were predicted to significantly increase as a function of bout length. **G**) Bout entropy normalized to bout length is significantly lower in KOs compared to WTs (p=6.53×10^−5^, W=26,904). Boxplots visualize the upper 75^th^ and lower 25^th^ percentiles of data with median values represented by horizonal bars. **H**) The rate of information gain in acortical KOs call bouts is lower, compared to WTs, and more similar to theoretical rate of information gain. **I**) Histogram comparing the distribution of WT (blue), KO (red), and theoretical (gray) normalized bout entropy. All distributions were significantly different (WT-KO: p=7.11×10^−5^, W=40,540; WT-Theoretical: p<2.2×10^−16^, W=285,778; KO-Theoretical: p p<2.2×10^−16^, W=157,816). Shaded area in, **F** and **H,** represent 95% confidence intervals. Statistical analysis of bout features computed using two-sided Wilcox rank sum test with continuity correction to evaluate effect of genotype on bout characteristics. ‘n.s.’, not significant; ‘*’, p≤0.05; ‘**’, p≤0.01; ‘***’, p≤0.001.

We then analyzed the amount of informational complexity contained within each call bout by computing Shannon’s entropy, a theoretical measure of information content, based on the relative frequency of syllable types within each bout. The average amount of entropy within call bouts was not significantly different between genotypes (p=0.530, W=32,633) (**Figure 3E**), an unexpected observation given that mutants had longer call bouts and the probability of incorporating new/additional syllable types would likely increase. To quantify this expectation, we computed the theoretical relationship between bout length and Shannon’s entropy using a re-sampled distribution of call bouts with sampling weights reflecting the proportion of syllable types produced by wildtype mice (**Figure 1B**). As expected, we found a significant and positive relationship between call bout length and entropy: bout entropy increased by 0.0068 units per additional syllable in each bout (**Figure 3F**). By computing the amount information content adjusted to bout length, termed ‘normalized entropy’, we find that acortical pups exhibit significant reductions in call bout complexity, compared to controls (p=7.12×10^−5^, W=40,540; **Figure 3G**). Moreover, the rate of information gain in acortical KOs is only 0.62x higher than expected by the null distribution of randomly re-sampled calls, whereas the rate of information gain among wildtypes was 3.3x higher than the theoretical rate of information gain (**Figure 3H**). The distribution of normalized entropy in acortical KOs is also significantly shifted more toward the theoretical null distribution (**Figure 3I**). Acortical pups appear capable generating isolation-induced call bouts, yet the bouts emitted by mutants have reduced temporal variation and informational complexity compared to controls.

### Impact of acortical and wildtype isolation calls on behavior of female dams

Pup USVs are an important part of the maternal-offspring interaction, as they allow pups to communicate internal states^16–18^. Our acoustic analyses revealed differences in the acoustic and temporal profile of the WT and acortical KO mice. We tested if mothers could detect differences in isolation call bouts by performing a modified version of the three-chamber task^19^. For this experiment, we generated female C57BL/6J dams with litters that age-match (P9) the original sample reported in Hammerschmidt et al (2015). After a 10-minute acclimation period, a composite recording of WT and acortical KO pups were simultaneously broadcasted from speakers located in opposite chambers of the three-chamber arena (**Figure 4A** and **4B**). The side of the chamber emitted the WT and KO recording was randomized. We then used Deep Lab Cut (DLC)^20^ to track the position of each female mouse’s body during the experiment and to measure the number of video frames that a dam resided in either chamber with a speaker (**Figure S2A**). We compared the difference in the percentage of frames each mouse spent in the playback chambers between the acclimation and playback periods to account of mouse-specific side preferences (see Methods section for details). When comparing the difference in percent residence between acclimation and playback, we found that during the first playback session, female dams increase the amount of time spent in the chamber with acortical KO playbacks compared to controls (**Figure 4D**; **Figure S3**). During the second playback, this proportional change in residence was non-significant (**Figure S2A**) with mice exhibiting no significant differences in residency location between the acclimation and playback periods (**Figure 4D**; **Figure S2B**). These results are consistent with previous studies showing rapid habituation within three-chamber environments with administration of sensory cues^21^. We found a significant difference in the residency shift between the two playback periods (**Figure 4D**), which further supports the conclusion that female preferences to search in the vicinity of mutant vocalizations decreased with the amount of habitation.

**FIGURE 4.**
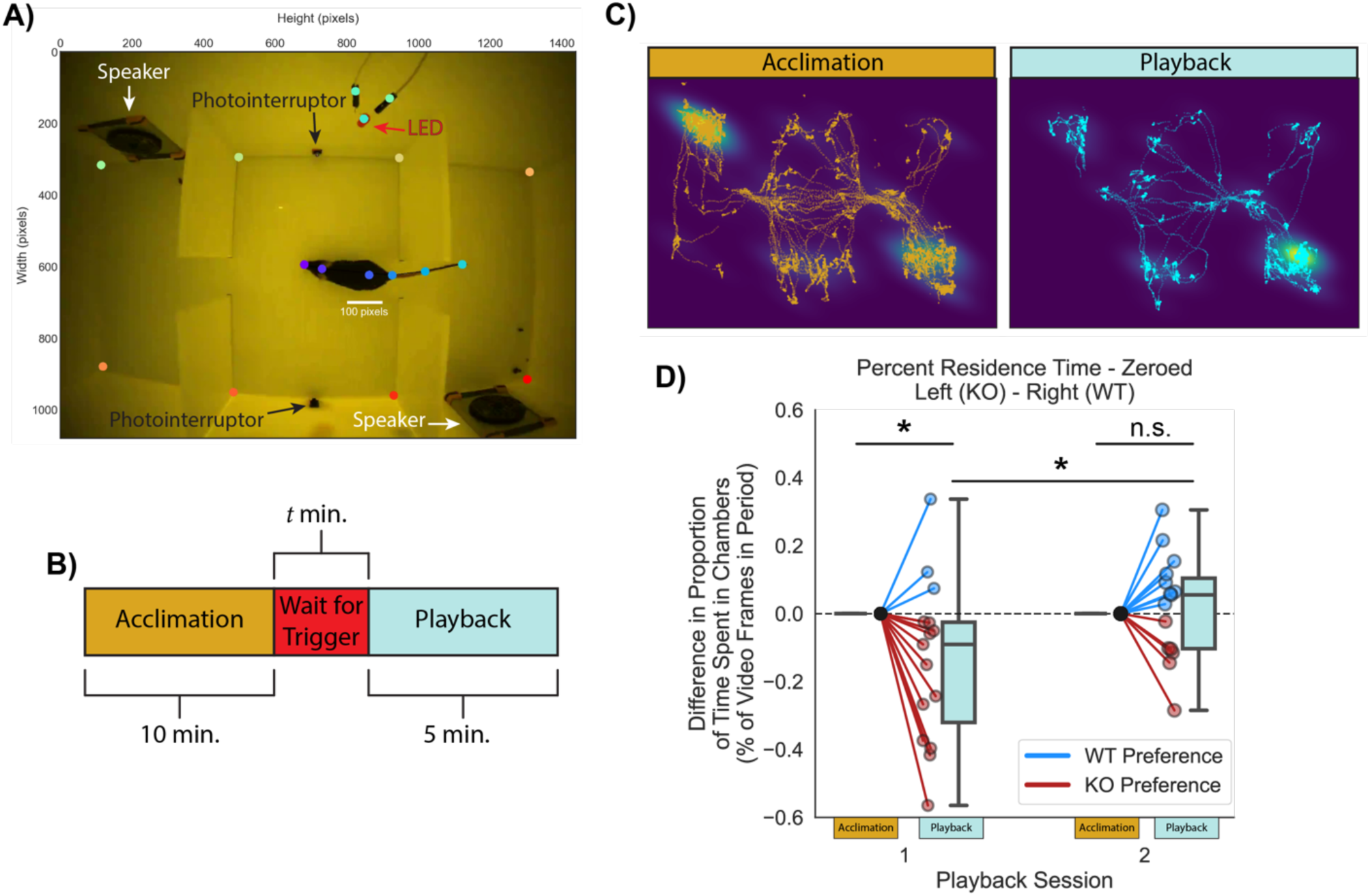
Playback of acortical pup vocalizations preferentially induce approach behavior by female dams. **A**) Single frame image of a female mouse in the three-chamber playback arena. Colored dots are the key points predicted from the trained DLC model. Scale bar is 100 pixels. **B**) Schematic of playback experiment timeline (see Methods for details). **C**) Example of cumulative traces of the tracked body position for a female throughout each period of the playback experiment. Traces overlayed on kernel density estimate (KDE) of each position across all frames for a given period. Regions of the highest density correspond to the location of each speaker. **D**) Boxplot and paired plots representing the change in each female mouse’s residence during the playback period as compared to the acclimation period. During the post-acclimation period of playback session 1, female dams spent significantly more time in the chamber with KO playbacks (red), compared to the chamber with WT playbacks (blue) (p=0.0364, df= 14, t=2.31). No differences were detected in residency during playback session 2 (p=0.602, df = 14, t=-0.533). Dam residency between playback session 1 and was significantly different (p=0.0336, df = 14, t=-2.36). Values are normalized to the percent duration (in frames) during the acclimation period for each mouse during each playback experiment. ‘n.s.’, not significant; ‘*’, p≤0.05.

## Discussion

In this study, we re-analyzed ultrasonic vocalizations of acortical mouse pups originally reported to be indistinguishable from wildtype control mice^10^. We extend findings from other re-analysis of vocal behavior in adult acortical mice by investigating the effects of specific acoustic features, analysis of temporal and informational dynamics, and playback experiments focused on assessment of vocalization-induced changes on maternal behavior^22^. We find that specific acoustic, temporal, and informational features help distinguish the vocal behavior of acortical mouse pups from wildtype pups. Our study reports several key findings: first, the vocal repertoires of acortical and wildtype pups differ on a subset of acoustic features; second, acortical pups produce vocalizations that are lower in pitch, louder in amplitude, and longer in duration compared to wildtype mice; third, isolation-induced call bouts of acortical pups have lower temporal variability and reduced informational complexity compared to controls that more closely resemble random chance; and fourth, females dams preferentially approach playbacks of isolation-induced calls emitted by acortical pups. Altogether, we find that mouse pups lacking significant areas of the cerebral cortex develop abnormal USVs.

### Interpretation of acoustic differences

The acoustic features of acortical pup vocalizations are consistent with stress-induced vocal responses. For example, maternal deprivation has been shown to induce the production of USVs with longer durations and lower frequencies compared to pups with lower or no maternal deprivation^16^. Observations of maternal behavior during our playback experiments are also consistent with previous studies that showed maternal dams preferentially approach pups producing USVs with higher amplitudes^23^. Similar to previous playback studies, the female mice in our experiment acclimated to the playbacks upon a second exposure^21^. Relative to the vocal features of wildtype pups, we hypothesize that the differences in acoustic features observed in acortical pups are consistent with higher levels of distress potentially associated with cortical dysregulation.

### Cortical control of vocal communication

The role of the cerebral cortex in mouse vocal communication remains contested with implications for experimental studies of vocal learning neural circuitry. In all vertebrates that vocalize and studied to date, subcortical circuits in the mid- and hindbrain serve to gate vocal production through premotor central pattern generations^11,12,24^. These brainstem circuits appear to be shared across all vertebrates^25,26^. On the other hand, classic dogma held that only in vocal learning species is the primary motor cortex (M1) required for proper vocal control and development^27,28^. Work from multiple groups report roles of different regions of M1 in modulating vocal behavior and flexibility in rodents^5,8^. The results of our re-analyses, paired with those from Ivanenko et al (2020), further demonstrate that, although the cortex is not required, or even the main contributor to vocal production in mice, its absence impacts the ability of mice to produce neurotypical vocalizations. Our results provide evidence that the cerebral cortex in non-vocal learning mammals helps to regulate several acoustic, temporal, and informational features of vocal behavior. Our playback study reinforces these findings, as the vocalizations are not only acoustically distinct between mutant and wildtype pups, but female dams changed their behavior in response to the vocalizations of pups lacking a cerebral cortex. Notably, even in vocal learning species, lesions to the motor cortex impact learned vocalizations, but these animals are still capable of producing innate vocalizations^5^. Thus, the rudimentary control of vocal behavior in mice may reflect their position on the vocal learning continuum, emphasizing their utility as a model for studying fundamental features of vocal behavior and evolutionary origins of speech and vocal learning.

### Comparison to previous analysis of vocal behavior of acortical pups

Why were the acoustic differences reported in our study not detected in the original Hammerschmidt et al (2015) study? We discuss several potential explanations. First, in our analysis of the recordings, generously provided by Kurt Hammerschmidt (18 total recordings; 8 WT, 10 KO), we detected more robust levels of vocal activity (number of syllables). Our USV extraction algorithm (MSA2) identified notably more individual USVs compared to the original study (133 USVs/min verse ~ 50 USV/min). For instance, the minimum level of vocal activity reported in our study included a mouse who emitted 261 USVs during the four minute recording session (65.3 USVs/min). By comparison, 12 out of 28 pups in the original Hammerschmidt et al (2015) study were reported to have vocal activities below 50 USVs/min, with ~8 pups emitting fewer, if any, USVs during the recording session. Detecting comparatively fewer USVs could mask statistical differences in acoustic features relative to larger vocalization dataset. Detection differences between algorithms are attributable to differences in computational methods and the effects of parameter specifications during extraction (e.g. settings for duration minimums, merge thresholds, etc.). Our extraction protocol utilized parameters that were previously identified as optimal for detection based on comparison to ground truths of manually labeled USVs^15^. Second, we focused analysis on statistical differences between genotypes based on the per animal average of each acoustic feature in order to avoid the implications of non-Gaussian distributions inherent to each acoustic feature (e.g. frequency bandwidth exhibits a bimodal distribution, while amplitude is exponential in nature) on linear or generalized models that test for statistical differences at the level of individual USVs. Third, many of the acoustic differences reported in our study that distinguished acortical KO from wildtype vocalizations (e.g. USV duration, mean frequency, amplitude) exhibit similar trends in data found in Hammerschmidt et al. (2015), albeit, they are not reported as statistically significant when analyzed at the level of individual USVs. As an additional line of support, maternal dams also appear to detect acoustic, temporal, or informational differences in the vocalizations of acortical pups compared to wildtypes.

In conclusion, we offer multiple lines of evidence to support the claim that mice lacking a cerebral cortex develop abnormal vocal behavior, which are detectable by dams with age-matched litters. Our evidence adds to the expanding literature on cortical control of vocalization and the neural foundations of vocal learning.

## Material and Methods

### Extraction of USVs and Vocal Features

#### Data

The ultrasonic sound recordings of mouse pup vocal behavior and metadata (genotypes, duration) originally reported in^10^, were generously provided by Kurt Hammerschmidt. The data included 18 WAV. format recordings of four minutes each, including wildtype (WT), *Esco2^+/+^;Emx1-Cre* (n=8), and acortical mutant knockouts (KO), *Esco2^fl/fl^;Emx1-Cre* (n=10), collected at postnatal day 9.

#### Algorithm

We used the Python implementation of MSA2 to generate sonograms from each WAV. file with the following parameters: purity bandwidth=1kHz, temporal resolution=2, snr threshold=10dB, purity threshold=0.5, discontinuity threshold=2, high pass frequency threshold=40kHz, duration threshold=6ms, merge threshold=15ms, frequency maximum=110kHz, frequency minimum=35kHz). Individual USVs were classified in syllable types based on pitch jumps: ‘s’ simple with no pitch jumps, ‘u’ up-jumps, ‘d’ down-jumps, and ‘M’ multi-jumps.

#### Statistical analysis

Numerical analysis was conducted in RStudio (Version 1.1.463) with R (4.0.3). Cluster analyses conducted with R packages *FactorMineR*, *factoextra*, *parameters* on default settings with scaled units for all acoustic features. The first two principal components were used for downstream analysis with maximum loading threshold and no rotation. The mean value acoustic features were calculated per animal per recording session. Data which failed the Shapiro-Wilk test for normality was evaluated using the Wilcoxon rank sum exact test. Statistical comparison of syllable types used linear models with genotype and syllable type as predictors. Main effects were assessed with Type III Analysis of Variance (ANOVAs) tables using the Satterthwaite’s method. Post hoc comparisons were computed with *emmeans* and Tukey adjustment for multiple test correction.

#### Analysis of call bouts

Sequences of five or more USVs separated by less than 250 ms of silence were grouped into call bouts. Bout acoustic features were calculated by averaging across all individual USVs within the bout. Additionally, we computed Shannon’s entropy using the *DescTools* package as a theoretical measure of information content. The amount of entropy (H) represented the minimum bits of information needed to encode the frequency of syllables types present in a call bout and was defined as: 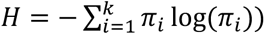, where *π* is the probability of syllable type *i* being present in a sequence of USVs for *k* syllable types, within a call bout. The null distribution of entropies for bouts of varying lengths were based on theoretical expectations of syllable frequences determine from randomly re-sampled call bouts at varying length (n=1,000 bootstrapped replicates).

### Playback Experiments

#### Animals

All original experiments in the present study were approved by the Institutional Animal Care and Use Committee (IACUC) at The Rockefeller University.

#### Female dams

In order to determine if wildtype mice could perceive differences between the normal and acortical pup vocalizations, we used female dams with litters that age-matched the animals used in the original study. Wildtype dams with the same genetic background as the original study were used (C57BL/6J). Playback experiments were scheduled to occur when dams had litters at P9.

#### Playback Files

Five minute composite sounds recordings were generated in Avisoft-SASLab Pro for each genotype (WT, KO) represented in the playback experiments. Composites were created by extracting one continuous minute of ultrasonic recording that contained a sequence of USVs. Five different source files were used to generate the composite file for each genotype. This approach accounted for the known inter-individual variability across animals and enabled us to account for varying levels of vocal production in the original data, which were determined to be approximately equivalent in the composite files.

#### Playback Experimental Design and Analysis

We constructed a three-chamber arena using cast acrylic (8505K744, McMaster-Carr), similar to Tajima et al. (2025). A camera (Firefly-S, FFY-U3-16S2C-S: 1.6 MP, 60 FPS, Sony IMX296, Color, FLIR) was placed in the center of the arena’s ceiling to allow us to track the mice’s position. In the chambers at each end, we placed ultrasonic playback speakers (Vifa ultrasonic speakers, Avisoft Bioacoustics, Berlin, Germany) on the wall facing in opposite directions (**Figure 4A**). This reduced the likelihood that the mice could hear the speaker from one chamber while in the opposite chamber. We determined the playback volume of each speaker by suspending an ultrasonic microphone (CM16/CMPA, Avisoft Bioacoustics) in the central chamber of the arena and observing the real-time spectrogram of the speakers using Avisoft-RECORDER. We lowered the volume of the speakers until neither speaker could be seen in the microphone’s spectrogram.

Each period of the experiments was controlled by a microcontroller (Feather M0, Adafruit, New York, NY). The acclimation period of the experiment lasted 10 minutes. After 10 minutes, the microcontroller waited for the mouse to pass between a set of photo interrupters (SEN-09299, SparkFun, Boulder, CO) located in the middle of the central chamber of the arena **Figure 4A**). This mechanism diminished the potential bias of having the mice already located in either playback chamber when the playbacks commenced. When the playbacks were initiated, an LED in the camera’s field of view was illuminated allowing us to determine when the playbacks started in our video analysis.

#### Analysis

To determine the experimental period from the video files, we created a custom pipeline in Bonsai^29^ that determined when the LED was illuminated, and provided a .csv that we could align to our position tracking frames (see below) to determine when mice were in the acclimation or playback periods. We tracked the animals’ position in the arena using Deep Lab Cut^30^. We trained a DLC resnet model to track five points on the mouse (**Figure S3A**), eight internal corners of the arena, and the LED. We decided to use the ‘body’ point on the mouse for our analyses based on the following considerations: 1) Based on DLC’s “likelihood” score for each point at each frame, the ‘body’ point had the highest proportion high-confidence points (>0.75) (**Figure S3B** and **S3C**). 2) We assessed the quality of each point’s tracking by determining the Euclidian distance (in pixels) of each point from one frame to the next (frame-to-frame point distance). Lower distances indicate greater continuity between frames and fewer ‘jumps’ in the tracking which are indicative of poor tracking performance or visual occlusions. We determined the ‘body’ point has the lowest frame-to-frame distances (**Figure S3B** and **S3D**).

Using the DLC tracked corners, we defined the borders of the left, right, and central chambers. Due to the jitter present in the DLC tracking output, we defined the coordinates of each corner by averaging the tracked position of the corner marker for each video. For each period of the experiment, we then determined the percentage of video frames that the animal spent in each playback chamber of the arena (left or right) as a binary state based on the location of the ‘body’ tracking point. (e.g. in a given frame, if the ‘body’ point was in a playback chamber but its ‘nose’ was in the central area the animal was determined to be in the playback chamber).

To standardize our analyses across our balanced playbacks, we assigned all KO playbacks as being in the left chamber, and all WT playbacks as being in the right chamber. To do this, we reversed the x-coordinate positions in our tracking for playbacks where the KO was originally assigned to the right. For each experimental period we calculated the ratio between left and right chamber residence (residence ratio). We determined that a mouse spent more or fewer video frames in a chamber based on the difference between the residence ratios of the acclimation and playback periods. A negative value means the mouse spent more time on the side with KO playbacks, and a positive value means the mouse spent more time on the side with the WT playbacks.

To perform our analyses we used custom Python code (Python 3.7) and the following freely available libraries: pandas, numpy, matplotlib, and seaborn.

## Acknowledgements

We thank Kurt Hammerschmidt for generously providing data of Esco2;emx-1-cre pup vocalizations and Peter F. Schade for assistance with MSA2, and other members of the Jarvis lab for comments and suggestions on this project.

## Author Contributions

J.L.B. conceived the study. J.L.B. and C.D.M.V performed experiments and analyzed data. J.L.B., C.D.M.V, and E.D.J. wrote the manuscript.

## Data Availability Statement

Recordings of *Esco2;Emx-1-Cre* mouse pup vocalizations originally obtained from Dr. Kurt Hammerschmidt. Other data and code for analysis available upon reasonable request.

## Supplemental Figures

**SUPPLEMENTAL FIGURE 1.**
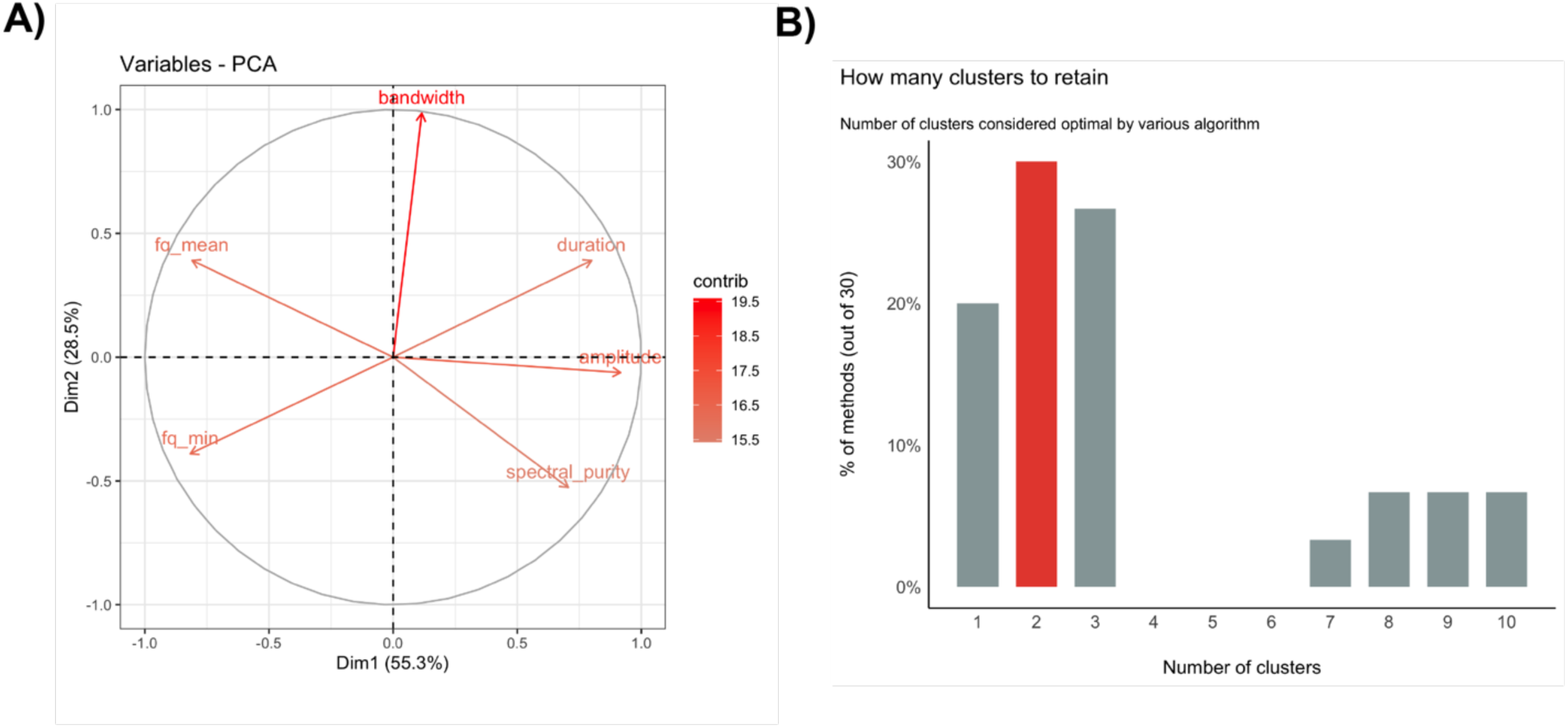
Principal component analysis of vocal features from wildtype and acortical mouse pups. **A**) Visualization of PCA vectors (cos2) for subset of vocal features with correlations less than 0.8 and projected into the first two dimensions. **B**) Histogram summarizing the consensus number of recommended clusters (red) based on the results of a battery of cluster optimization algorithms, including elbow, gap, silhouette, dbscan, hclust, which compute the optimal number of clusters. Visualization of PCA outputs performed in R Studio package *factoextra* and cluster optimization computed using n_*clusters* package on standardized data.

**SUPPLEMENTAL FIGURE 2.**
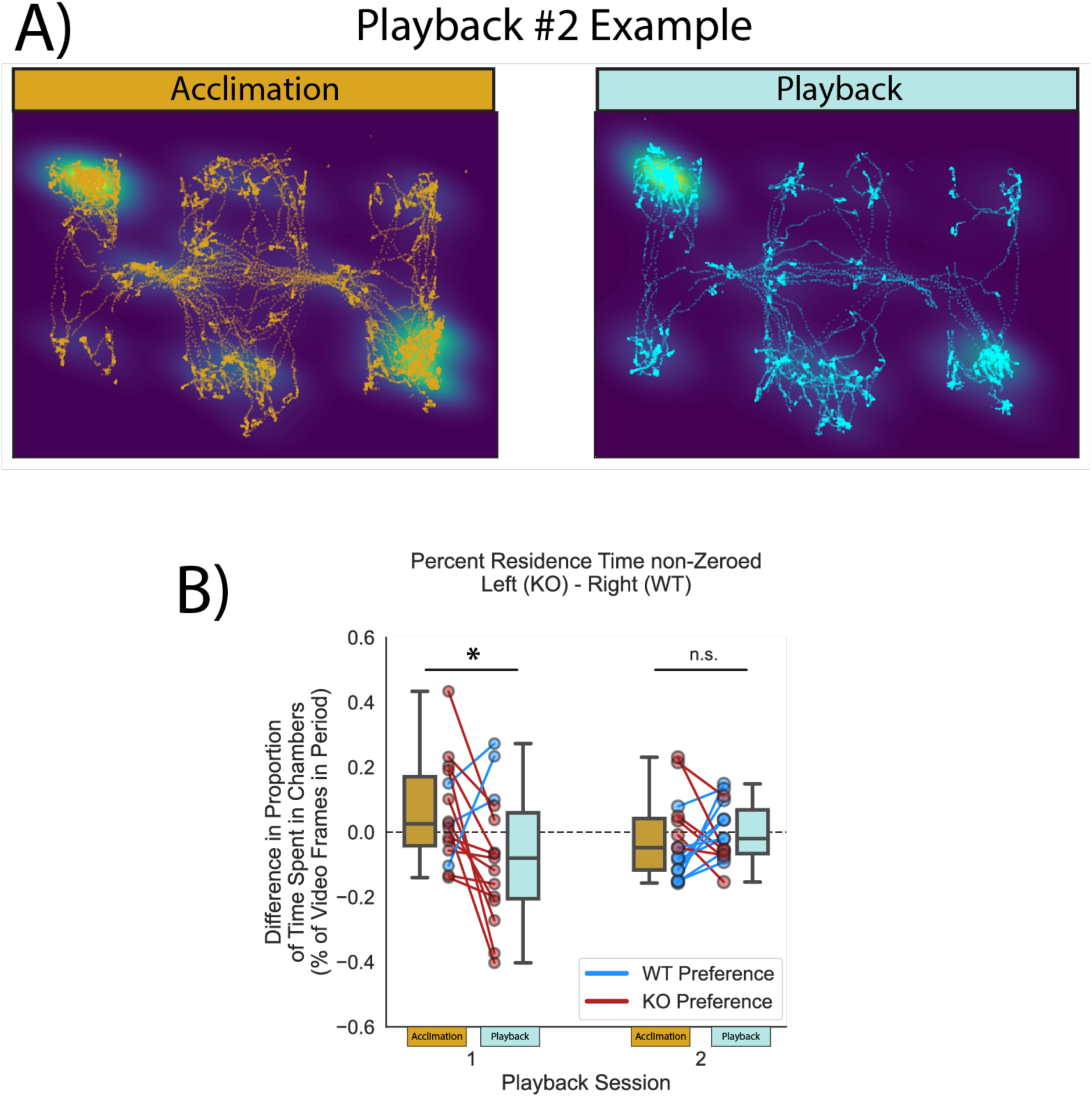
Playback preference of acortical female mice is not maintained during a second playback session. **A**) Example of cumulative traces of the tracked body position for a female throughout each period of the second playback experiment from the same mouse as (Figure 4C). Traces overlayed on kernel density estimate (KDE) of each position across all frames for a given period. Regions of the highest density correspond to the location of each speaker. **B**) Boxplot and paired plots representing the non-normalized change in each female mouse’s residence during the playback period as compared to the acclimation period. During the post-acclimation period of playback session 1, female dams spent significantly more time in the chamber with KO playbacks (red), compared to the chamber with WT playbacks (blue) (p=0.0364, df=14, t= 2.31). No differences in residency were observed during playback session 2 (p=0.602, df=14, t=-0.533). ‘n.s.’, not significant; ‘*’, p≤0.05.

**SUPPLEMENTAL FIGURE 3.**
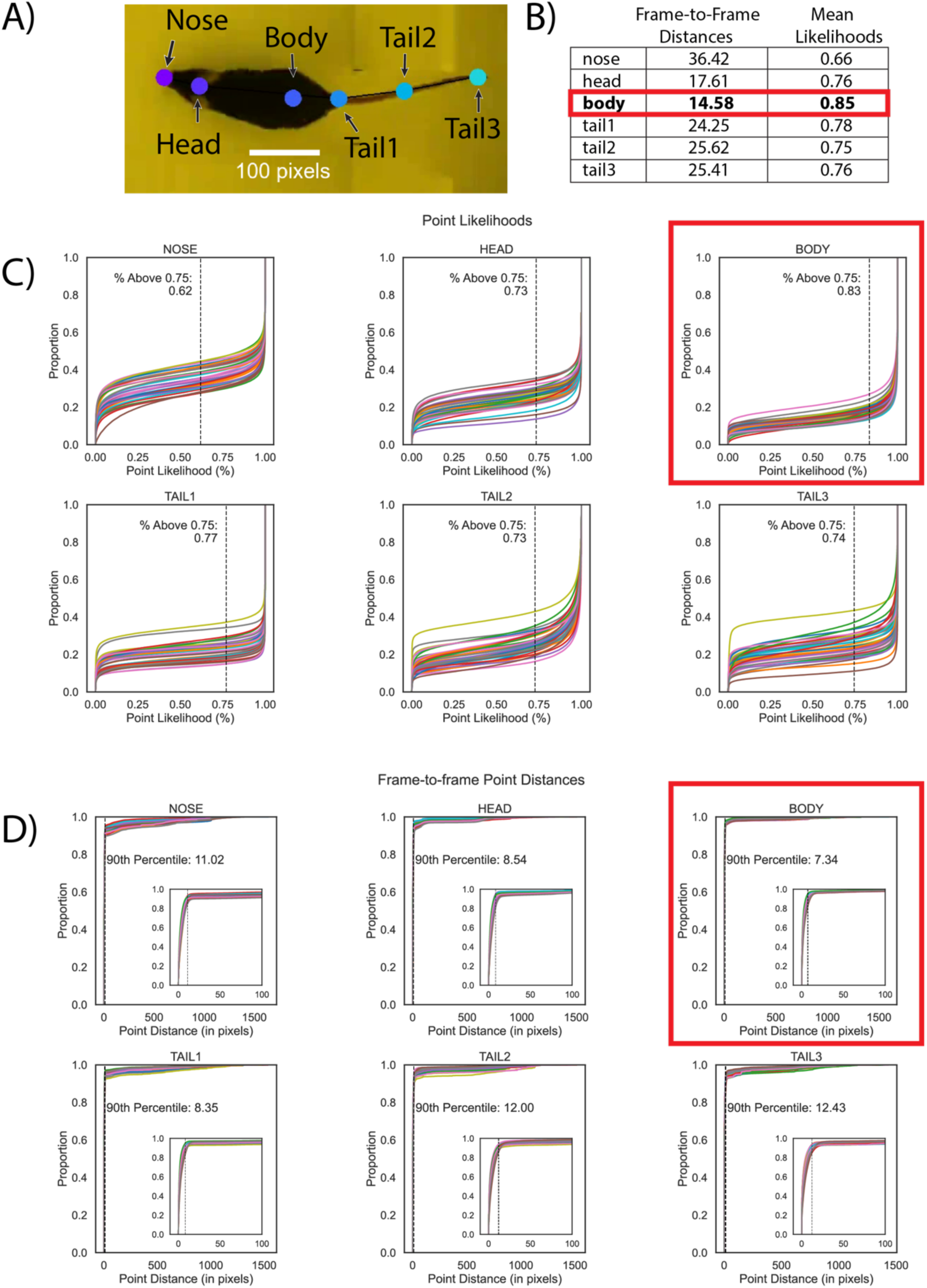
Metrics for each point tracked on mice in playback videos. **A**) Example image of point skeleton and labels on a mouse that has been tracked. **B**) Table summarizing mean values of frame-to-frame distance and mean likelihood scores showing the body point (boxed in red) had the best performance overall. **C**) Empirical cumulative distribution plot (ECDF) of all likelihood values for each point. Each line in a plot represents the values of each video for a given point. Body (red box) had the highest number of points above 0.75 likelihood score and had the narrowest distribution suggesting greater consistency in tracking. **D**) ECDF of point-to-point distances across frames for each point. Each line in a plot represents values for each video for a given point. Overall, body (boxed in red) had the lowest distance at the 90^th^ percentile, suggesting more consistent tracking of that point between frames.

## References

1. Jarvis, E. D. Evolution of vocal learning and spoken language. Science 366, 50–54 (2019).

2. Petkov, C. I. & Jarvis, E. D. Birds, primates, and spoken language origins: behavioral phenotypes and neurobiological substrates. Front. Evol. Neurosci. 4, (2012).

3. Wirthlin, M. et al. A Modular Approach to Vocal Learning: Disentangling the Diversity of a Complex Behavioral Trait. Neuron 104, 87–99 (2019).

4. Fitch, W. T. Empirical approaches to the study of language evolution. Psychon Bull Rev 24, 3–33 (2017).

5. Arriaga, G., Zhou, E. P. & Jarvis, E. D. Of Mice, Birds, and Men: The Mouse Ultrasonic Song System Has Some Features Similar to Humans and Song-Learning Birds. PLoS ONE 7, e46610 (2012).

6. Vargas, C. D. M. et al. A Functional and Non-Homuncular Representation of the Larynx in the Primary Motor Cortex of Mice, a Vocal Non-Learner. Preprint at 10.1101/2024.02.05.579004 (2024).

7. Van Daele, D. J. & Cassell, M. D. Multiple forebrain systems converge on motor neurons innervating the thyroarytenoid muscle. Neuroscience 162, 501–524 (2009).

8. Banerjee, A., Chen, F., Druckmann, S. & Long, M. A. Temporal scaling of motor cortical dynamics reveals hierarchical control of vocal production. Nat Neurosci 27, 527–535 (2024).

9. Isko, E. C. et al. Selective expansion of motor cortical projections in the evolution of vocal novelty. Preprint at 10.1101/2024.09.13.612752 (2024).

10. Hammerschmidt, K., Whelan, G., Eichele, G. & Fischer, J. Mice lacking the cerebral cortex develop normal song: Insights into the foundations of vocal learning. Sci Rep 5, 8808 (2015).

11. Park, J. et al. Brainstem control of vocalization and its coordination with respiration. (2024).

12. Tschida, K. A Specialized Neural Circuit Gates Social Vocalizations in the Mouse.

13. Holy, T. E. & Guo, Z. Ultrasonic Songs of Male Mice. PLoS Biol 3, e386 (2005).

14. Chabout, J., Sarkar, A., Dunson, D. B. & Jarvis, E. D. Male mice song syntax depends on social contexts and influences female preferences. Front. Behav. Neurosci. 9, (2015).

15. Stoumpou, V. et al. Analysis of Mouse Vocal Communication (AMVOC): a deep, unsupervised method for rapid detection, analysis and classification of ultrasonic vocalisations. Bioacoustics 32, 199–229 (2023).

16. Yin, X. et al. Maternal Deprivation Influences Pup Ultrasonic Vocalizations of C57BL/6J Mice. PLoS ONE 11, e0160409 (2016).

17. Wöhr, M., Engelhardt, K. A., Seffer, D., Sungur, A. Ö. & Schwarting, R. K. W. Acoustic Communication in Rats: Effects of Social Experiences on Ultrasonic Vocalizations as Socio-affective Signals. in Social Behavior from Rodents to Humans (eds. Wöhr, M. & Krach, S.) vol. 30 67–89 (Springer International Publishing, Cham, 2015).

18. Premoli, M., Memo, M. & Bonini, S. Ultrasonic vocalizations in mice: relevance for ethologic and neurodevelopmental disorders studies. Neural Regen Res 16, 1158 (2021).

19. Tajima, Y. et al. A humanized NOVA1 splicing factor alters mouse vocal communications. Nat Commun 16, (2025).

20. Mathis, A. et al. DeepLabCut: markerless pose estimation of user-defined body parts with deep learning. Nat Neurosci 21, 1281–1289 (2018).

21. Hammerschmidt, K., Radyushkin, K., Ehrenreich, H. & Fischer, J. Female mice respond to male ultrasonic ‘songs’ with approach behaviour. Biol. Lett. 5, 589–592 (2009).

22. Ivanenko, A., Watkins, P., Van Gerven, M. A. J., Hammerschmidt, K. & Englitz, B. Classifying sex and strain from mouse ultrasonic vocalizations using deep learning. PLoS Comput Biol 16, e1007918 (2020).

23. Wöhr, M. et al. Effects of Genetic Background, Gender, and Early Environmental Factors on Isolation-Induced Ultrasonic Calling in Mouse Pups: An Embryo-Transfer Study. Behav Genet 38, 579–595 (2008).

24. Michael, V., et al. Circuit and synaptic organization of forebrain-to-midbrain pathways that promote and suppress vocalization. eLife 9, e63493 (2020).

25. Jürgens, U. The Neural Control of Vocalization in Mammals: A Review. Journal of Voice 23, 1–10 (2009).

26. Bass, A. H., Gilland, E. H. & Baker, R. Evolutionary Origins for Social Vocalization in a Vertebrate Hindbrain–Spinal Compartment. Science 321, 417–421 (2008).

27. Jürgens, U. Neural pathways underlying vocal control. Neuroscience & Biobehavioral Reviews 26, 235–258 (2002).

28. Fitch, W. T., Huber, L. & Bugnyar, T. Social Cognition and the Evolution of Language: Constructing Cognitive Phylogenies. Neuron 65, 795–814 (2010).

29. Lopes, G. et al. Bonsai: an event-based framework for processing and controlling data streams. Front. Neuroinform. 9, (2015).

30. Mathis, A. et al. DeepLabCut: markerless pose estimation of user-defined body parts with deep learning. Nat Neurosci 21, 1281–1289 (2018).

